# Structure basis for single-strand nucleic acid targeting by IscB and variants

**DOI:** 10.64898/2026.03.03.709405

**Authors:** Chengtao Xu, Qi Yang, Xiaolin Niu, Ailong Ke

**Affiliations:** Department of Molecular Biophysics and Biochemistry, Yale University, New Haven, CT 06511, U.S.A.

## Abstract

Transposon-encoded IscB has been established as the evolutionary ancestor of CRISPR-Cas9. This compact RNA-guided endonuclease has since been engineered for genome editing applications. We previously repurposed IscB and Cas9 as efficient RNA editors by removing their double-stranded DNA recognition module, the TAM/PAM-interacting domain. Here, we report four cryo-EM structures of IscB in complex with single-stranded nucleic acid (ssNA) targets to illuminate its mechanistic underpinnings. Structural analysis reveals that IscB initially facilitates formation of a 10-nt seed duplex with ssNA; however, further base-pairing is blocked by an alternatively positioned HNH nuclease that acts as a conformational roadblock. In this intermediate state, neither HNH nor RuvC is competent for target cleavage: the HNH domain is occluded by the roadblock configuration, while the RuvC active site is obstructed by the guide RNA. Only upon full duplex formation do additional base pairs between the guide RNA and ssNA dislodge the HNH roadblock, simultaneously exposing the RuvC nuclease active site. We propose that an analogous conformational checkpoint governs IscB activity during dsDNA target interrogation. Guided by these structural insights, we introduced mutations to either enhance ssNA binding or relieve the conformational checkpoint, both of which significantly improved RNA-targeting efficiency of IscB.

## Introduction

IscB is a compact RNA-guided endonuclease encoded by IS200/IS605 transposon elements. It is considered the evolutionary ancestor of the CRISPR-associated effector Cas9^1-3^. While only two-fifths the size of Cas9, IscB shares a similar domain architecture, featuring an arginine-rich bridge helix and an HNH endonuclease domain inserted into a split RuvC nuclease domain^4,5^. In Cas9, the bridge helix nucleates the assembly of a ribonucleoprotein (RNP) complex with two noncoding RNAs, CRISPR RNA (crRNA) and trans-activating CRISPR RNA (tracrRNA)^6-10^. Double-stranded DNA (dsDNA) is accommodated into Cas9 by the protospacer adjacent motif (PAM) interaction domain (PID) and unwound to form the so-called R-loop structure, where the target-strand (TS) DNA forms a heteroduplex with the guide portion of crRNA and the non-target-strand (NTS) DNA loops out. TS and NTS are subsequently cleaved by HNH and RuvC nucleases, respectively^11^. A large recognition (REC) lobe composed of multiple α-helical domains facilitates DNA unwinding and R-loop formation^9,11-16^. In IscB, this REC lobe is replaced by the scaffold portion of ωRNA (obligate mobile element-guided activity RNA)^3^, the functional equivalent of crRNA and tracrRNA. Target DNA recognition in IscB is initiated through TAM (target-adjacent motif) recognition, a mechanism that mirrors PAM recognition in Cas9^17,18^. R-loop formation and the DNA cleavage mechanism in IscB are mechanistically conserved as in Cas9^4,5,9,11-16^. IscB additionally encodes an N-terminal PLMP-motif-containing domain, which is important for RNP assembly^3^.

Following its discovery and structure determination^4,5,19,20^, IscB has been extensively engineered to improve its genome editing performance in mammalian cells^19-28^. For example, we improved the editing efficiency of *O. geu* IscB by thirty folds in human cells, by introducing an eight-amino-acid (8-aa) substitution to OgeuIscB^25^. Recently, we further repurposed IscB into ssDNA and ssRNA-targeting enzymes, by removing its TID domain. The resulting R-IscB variant is quite efficient in mediating mRNA knockdown, splicing perturbation, A-to-I editing, and trans-splicing^28^. Here we aimed to gain deeper mechanistic understanding of its ssNA targeting activity by determining high-resolution structures. In this study, we report the cryo-electron microscopy (cryo-EM) structures of the full-length IscB and the TID-less R-IscB, bearing their corresponding beneficial mutations and bound to ssDNA or ssRNA. We confirm that the beneficial mutations for dsDNA targeting do not alter IscB structure appreciably, therefore they likely function through improved nucleic acid contacts. Importantly, we found that RNA-guided ssNA recognition in both full-length and TID-less IscB proceed through a two-step process. IscB first promotes the formation of a 10-nt seed duplex between guide RNA (gRNA) and target ssNA; additional base-pairing was blocked by the alternatively docked HNH at this functional state. Neither HNH nor RuvC is capable of cleaving targets in the seed-duplex state. The HNH active site is oriented to the backside of IscB and RuvC active site is blocked by the side-tracked gRNA. In a second snapshot gRNA is fully base-paired with the DNA target, the HNH roadblock is dislodged to an alternative conformation, and the RuvC active site becomes exposed. Inspired by the structural observations, we introduced mutations into R-IscB to either favor ssNA binding or facilitate the seed-to-full duplex transition. A high proportion of variants show improved target-binding behaviors. Overall, this is a productive effort to improve R-IscB into a robust single-stranded genome editor.

## Results

### Cryo-EM reconstruction of ssDNA-bound IscB reveals two conformational states

We previously showed that the full-length OgeuIscB was also capable of ssNA binding, albeit it strongly preferred dsDNA targets. We traced the substrate preference to its TID domain, the deletion of which converted IscB into an exclusive ssNA binder^28^. To first understand how ssNA is accommodated in the full-length OgeuIscB, we incubated it with a guide-complementary ssDNA at 37 °C for 20 minutes before snap-freezing the sample for cryo-EM single-particle reconstruction (Figures 1A, 1B and S1). Two conformational states were resolved from this dataset (Figures 1C-E and S2). They differ significantly in the extent of the duplex between gRNA and target ssDNA (tDNA), the HNH nuclease location, and the RuvC status. As explained below, these structural differences are consistent with two sequential events in ssDNA binding, a seed-duplexed state (2.86 Å in structural resolution, Figures 1D and S2-3) corresponding to a partially bound ssDNA target, and a fully-duplexed state (2.84 Å, Figures 1E and S2) corresponding to a fully recognized ssDNA target. The former state is reconstructed from ∼30% of the total single particles whereas the latter ∼35%, no other distinct conformations can be reconstructed to high-resolution from the remaining ∼35% of the particles (Figure S2). These observations support our previous conclusion that IscB recognizes ssNA through seed-sequence recognition^28^, and further suggest that a kinetic bottleneck in the form of a conformational checkpoint governs the transition from the seed-duplexed to the fully-duplexed state.

**Figure 1.**
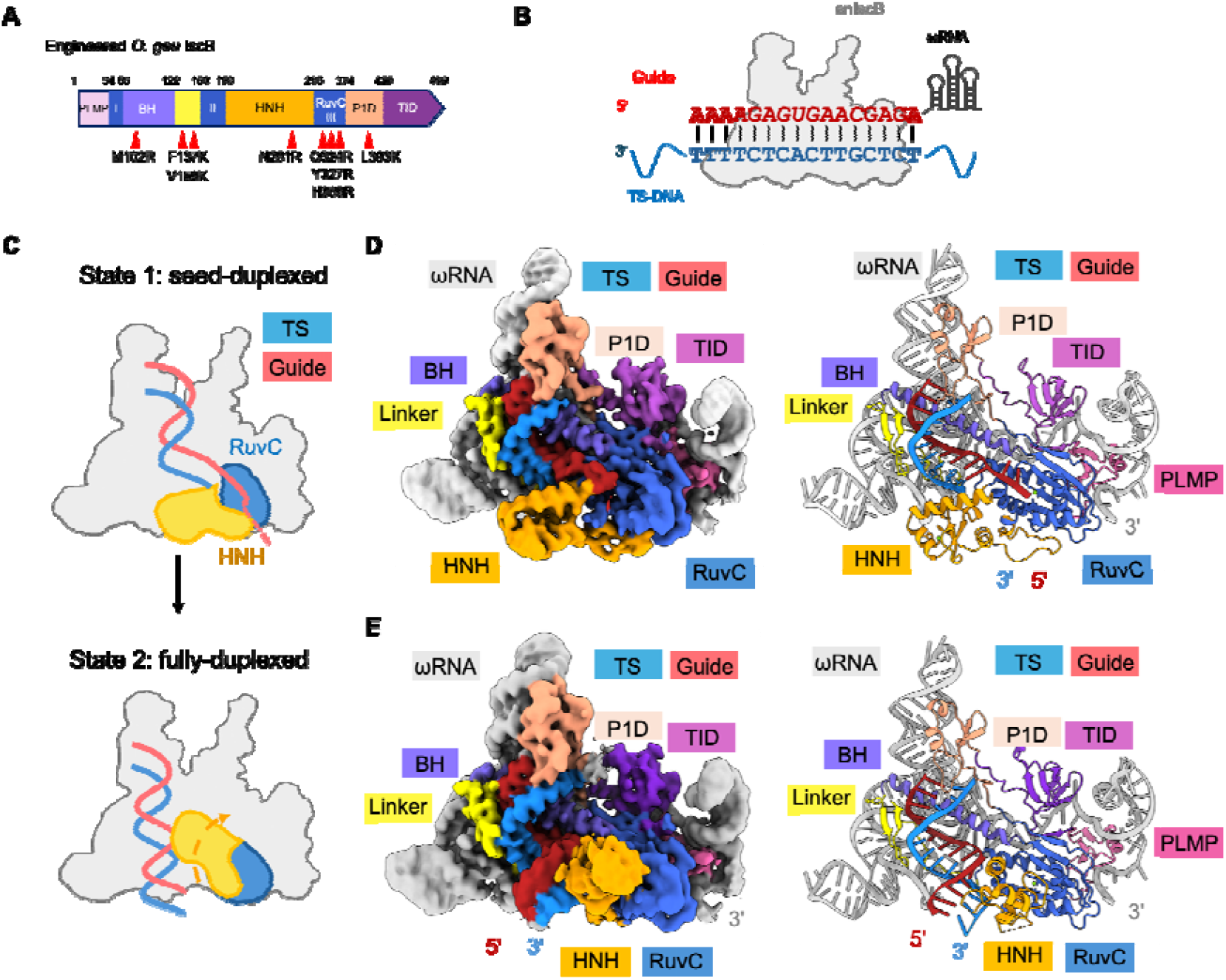
Cryo-EM reconstruction of enIscB boud to ssDNA revealed two conformational states, seed-duplexed and fully-duplexed. (A) Domain organization of enIscB^25^ with beneficial mutations improving genome editing in human cell editing marked. Color scheme is consistent throughout Figure 1. (B) Diagram of hybridization formed between guide RNA and target ssDNA. (C-E) Seed-duplex (top) and full-duplex state (bottom) illustrated in domain organization (C) and in cryo-EM densities their corresponding structural models (D-E). Note the differences in HNH domain location, guide RNA direction, and gRNA/tDNA length.

The previously determined dsDNA-bound IscB structure (PDB: 7UTN) docks seamlessly into our newly obtained cryo-EM map (Figure 2A). The TID domain of IscB adopts the same overall orientation even when TAM-proximal dsDNA is absent (Figures 2B and S4). While the overall conformation of the gRNA–tDNA heteroduplex is largely similar, the density corresponding to the first nucleotide of the heteroduplex is significantly weaker in the ssDNA-bound structures (Figure 2C). This base pair appears to be disfavored by the nearby P1D domain lip of IscB (K400–T406), whose bulky residues impose geometric distortion on the local base pairs. This distortion is avoided when IscB binds dsDNA, as the TAM-proximal dsDNA pulls the downstream TS DNA further away. We speculate that the P1D lip may bias IscB toward preferring dsDNA over ssNA.

**Figure 2.**
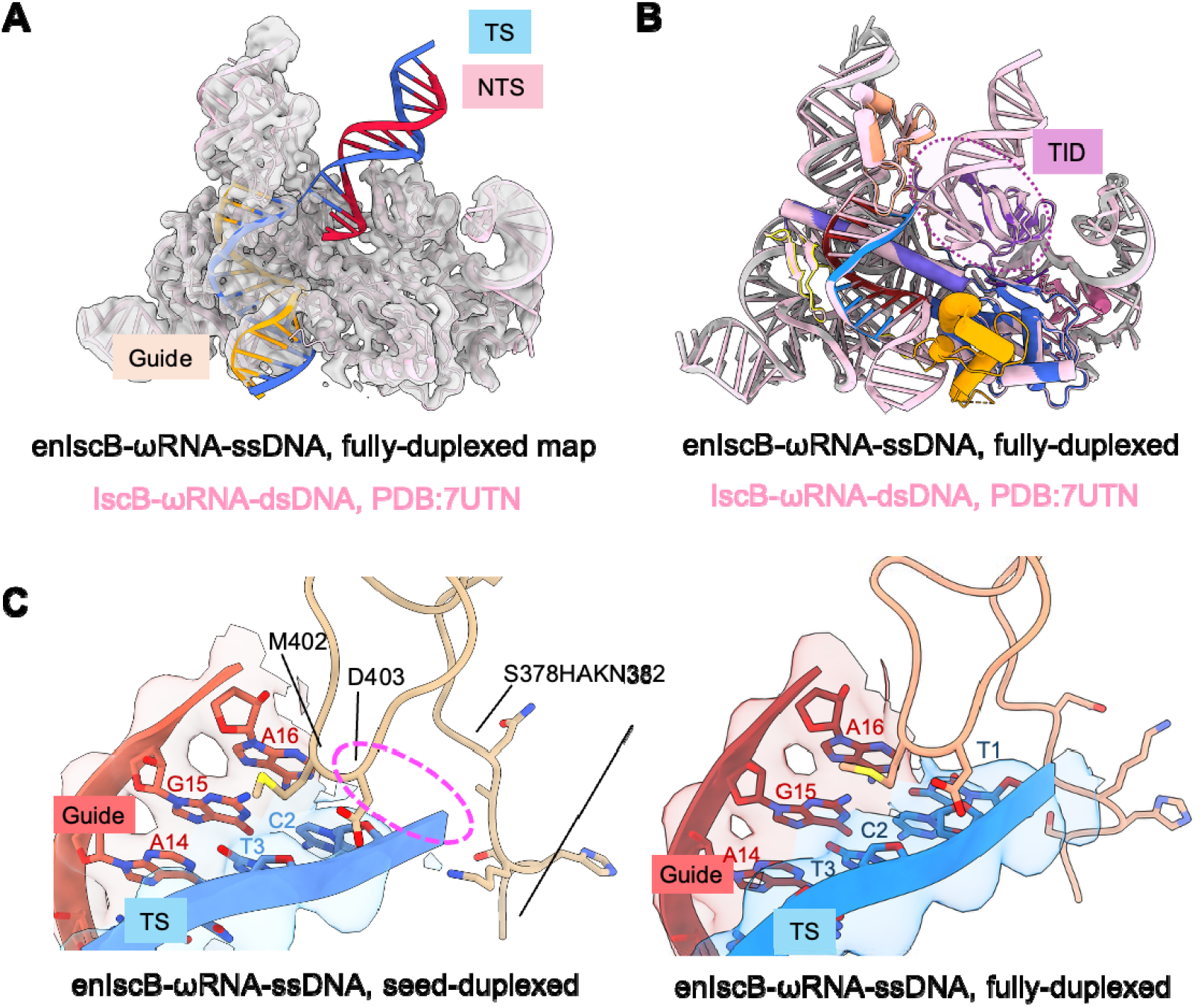
Structure comparison reveals similarity of IscB architecture between dsDNA-bound and ssDNA-bound state. (A) Previously reported IscB-ωRNA-dsDNA structure (PDB:7UTN) can be fit in the fully-duplexed enIscB-ωRNA-ssDNA density map (this study). (B) Overlay of fully-duplexed enIscB-ωRNA-ssDNA and IscB-ωRNA-dsDNA shows high similarity, including the organization of TID. (C) Left: electron density of the first nucleotide (T1) on the TS-DNA is missing in the seed-duplexed state; right: the first nucleotide can be identified in the fully-duplexed state.

### HNH roadblock and nuclease sequestration in the seed-duplexed state

The most distinctive structural feature of the seed-duplexed state is the alternatively packed HNH nuclease. Previous structures captured IscB only in the fully R-looped state, in which HNH is either invisible due to high mobility (PDB: 8CTL) or bound to the gRNA-tDNA heteroduplex in the pre-catalytic state (PDB: 8CSZ)^5^. Here we capture an intermediate state in which HNH serves as a roadblock, preventing propagation of the gRNA-tDNA heteroduplex (Figure 3A). HNH is stabilized in this conformation through contacts mediated by a linker connecting RuvC to HNH (N194–S221, Figure 3B). The function of this linker was previously poorly understood, and it was poorly defined in previous structures. Here we show that residues R195-M199 within this linker fold into a short helix that bridges an extensive network of hydrophobic contacts between HNH and RuvC (Figure 3C). In particular, F196 at the tip of the helix extends into a hydrophobic pocket in HNH formed by Y293, M228, N229, and L258. Beyond the short helix, the linker region makes two layers of contacts against RuvC (Figure 3D). The most noteworthy interactions include hydrophobic contacts from K205, V206, W209, and Y211 to the surface of RuvC.

**Figure 3.**
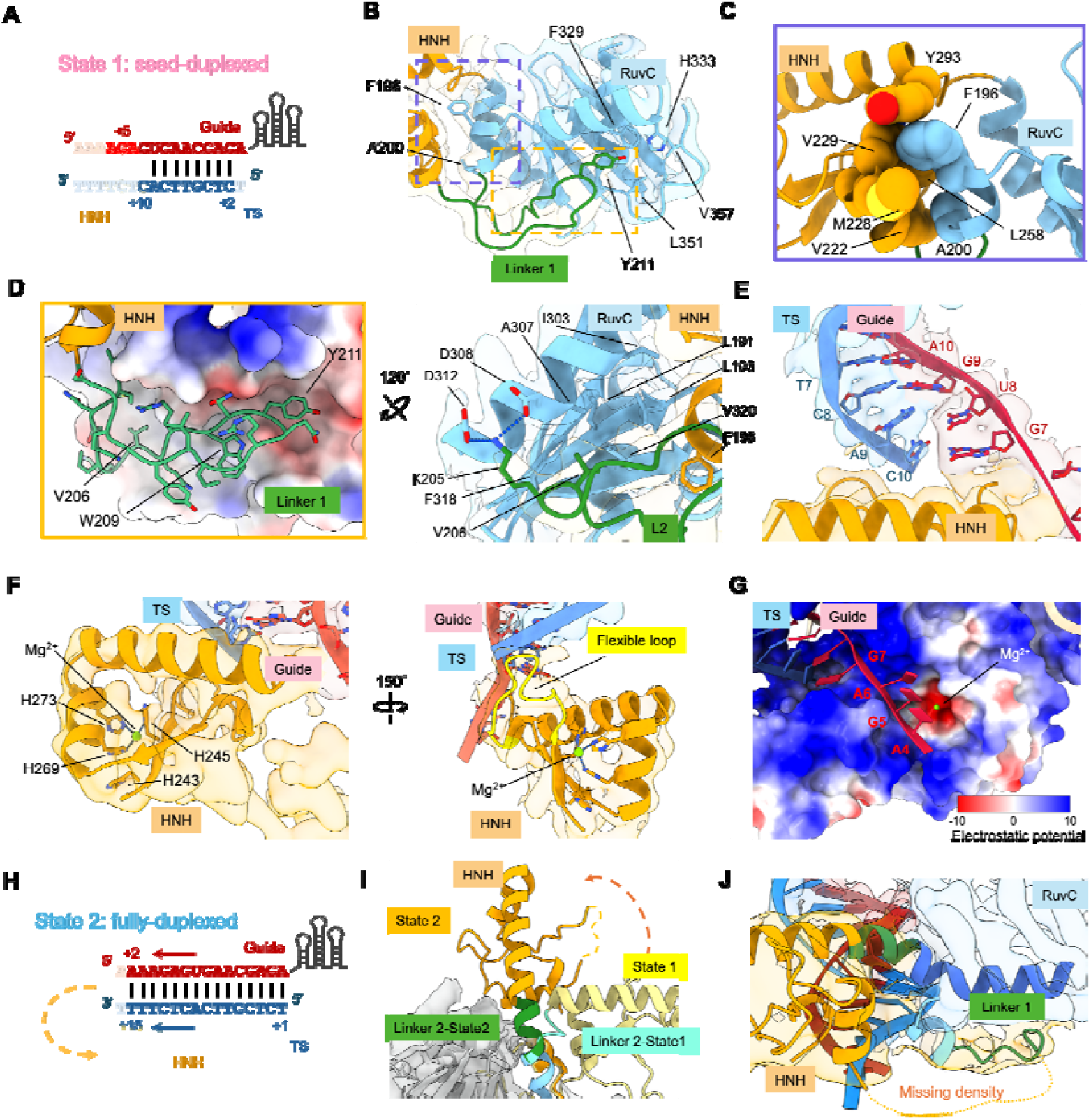
Structural differences between the seed-duplex and full-duplex states. (A) Extent of gRNA-tDNA base-pairing in the seed-duplexed state. (B) Flexible linker 1 on HNH shows extensive interactions with RuvC domain in the seed-duplexed state. The boxed regions are analyzed in zoom-in panels in (C-D). (C) Short helix (R195-M199) shows extensive hydrophobic contacts between HNH and RuvC in the shape of a ball-socket joint. (D) Flexible linker lays in the pocket at the bottom of RuvC domain. Two views depicting residues that form hydrophobic interactions and salt bridges with those in the RuvC domain. (E) Blockage of gRNA-tDNA extension in the seed-duplex state by the alternatively packed HNH domain. The Watson-Crick base-pairing of the last nucleotide C10 with G7 is disrupted due to steric clash. (F) Domain architecture and geometry at the catalytic core of HNH domain in the seed-duplexed state illustrated in two views. (G) Blockage of RuvC active site in the seed-duplex state by the detoured gRNA, as explained by the structural model superimposed with the electrostatic potential surface. (H) Extent of gRNA-tDNA base-pairing in the fully-duplexed state. (I) The reorganization of linker 2 induces the 90° rotation of HNH domain from the seed-duplexed to the fully-duplexed state. (J) The majority of linker 1 cannot be identified in the fully-duplexed state due to increased mobility.

Trapping HNH in the path of the gRNA leads to three significant functional consequences. First, it orients the HNH active site toward the backside of IscB, preventing it from cleaving partially base-paired DNA and RNA targets (Figure 3E). In this new location, the HNH active site is flanked by the P2 stem of ωRNA, which may restrict access of non-target nucleic acids for cleavage. Second, the first ten nucleotides of the gRNA follow the line of positive charges along the bridge helix until encountering the HNH roadblock (G7 to A16), base-pairing with the complementary nucleotides in ssDNA to form the seed duplex. The first and last ssDNA residues are poorly resolved in the duplex, suggesting that they are frequently unpaired (Figures 2C and 3F). Lastly, after being redirected by the HNH roadblock, the guide portion of ωRNA detours into the RuvC active site. A stretch of three nucleotides (A4 to A6) can be traced, and the density suggests that the gRNA traverses the RuvC active site uncleaved, possibly due to suboptimal orientation and geometry (Figure 3G). Although not cleaved, the gRNA physically occludes the RuvC active site, preventing it from accommodating ssDNA substrates for cleavage. Thus, neither HNH nor RuvC is capable of cleaving the RNA-guided target in the seed-duplexed state.

### Conformational rearrangement and nuclease activation in seed-to-full duplex transition

The fully-duplexed state depicts the aftermath of seed-to-full-duplex transition. The gRNA-tDNA heteroduplex extended to its full extent (15 bp modeled) and the overall conformation is similar to that depicted in the dsDNA-bound IscB structure (PDB: 7UTN, Cα r.m.s.d = 0.667Å, Figures 3H and S5). This is coupled with the disappearing of gRNA densities from the RuvC active site and the dislodging of the HNH roadblock. In the seed state, the last α-helix of HNH and the following α-helix in RuvC are separated by a 90° elbow in the form of a 4-aa linker (H294-A295-L296-S297). In the fully-duplexed state, the elbow straightens as the linker morphs into an imperfect α-helix, enabling the formation of a long, slightly curved α-helix, from HNH into RuvC (Figure 3I). Concurrently, the contacts between the HNH-to-RuvC linker and RuvC that were present in the seed-duplexed state are lost, and this linker becomes absent from the cryo-EM density map, suggesting it becomes disordered (Figure 3J). Notably, the HNH densities are only visible at low contour level, and the local resolution only allowed rigid-body docking of the HNH core structure (A221-H294). HNH is either highly mobile or occupies this conformation only transiently. These two possibilities are not mutually exclusive.

### ssRNA-bound R-IscB structures defines the impact of TID deletion

TID domain deletion was the key step in converting dsDNA-targeting IscB to ssNA-targeting R-IscB. To explore avenues for further improving its RNA-targeting activity, it is important to first characterize the structural impact of TID deletion. We therefore programmed R-IscB with an ssRNA target and determined its cryo-EM structure (Figures 4A, 4B, S1, and S6). As with full-length IscB, R-IscB adopted two conformational states, the seed-duplexed state and the fully-duplexed state, each resolved from approximately equal populations of single particles (Figures 4C–E and S6). This suggests that neither TID deletion nor substrate identity (ssRNA versus ssDNA) significantly alters the conformational switching-based target recognition mechanism. The two conformational states superimpose extremely well with their counterparts in full-length IscB (r.m.s.d. in the 0.5–0.6 Å range; Figure S5), further indicating that TID deletion does not significantly alter the overall IscB structure. Interestingly, the geometry of the gRNA–tRNA duplex is superimposable with the gRNA–tDNA heteroduplex observed in previously solved structures, despite the additional 2’-hydroxyl groups, and deviates from the standard A-form with significantly widened major grooves. This observation suggests that the duplex geometry is dictated by IscB contacts, including the line of positive charges along the bridge helix. Some local structural changes surrounding the deleted TID are noted. In an effort to ensure correct folding of the R-IscB RNP, we preserved the flexible linker from P1D to TID and fused it directly to the β-strand following TID, which zips to the edge of the β-sheet in RuvC. The R-IscB structure reveals, however, that neither the linker nor the β-strand is required for the structural integrity of R-IscB: while the entire P1D domain is well defined in the cryo-EM density, the linker beyond P1D (after Y433) and the fused β-strand are absent from the map (Figure 4F).

**Figure 4.**
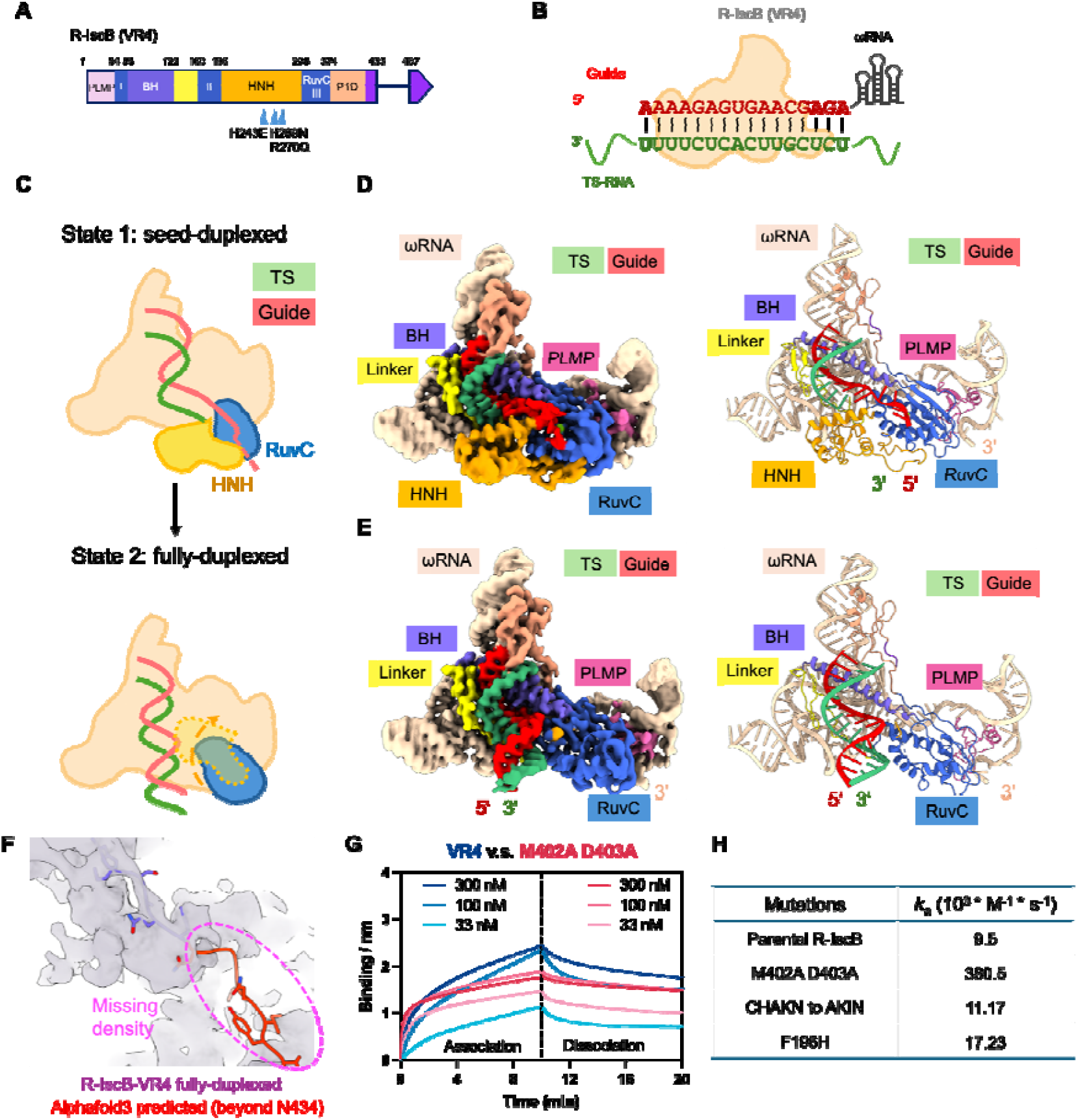
Cryo-EM reconstruction of R-IscB-VR4 bound to ssRNA revealed two conformational states, seed-duplex and full-duplex. (A) Domain organization of R-IscB-VR4^28^ with ssRNA-cleaving mutations marked. Color scheme is consistent throughout Figure 4. (B) Diagram of hybridization formed between guide RNA and target ssRNA. (C-E) Seed-duplex (top) and full-duplex state (bottom) illustrated in domain organization (C) and in cryo-EM densities their corresponding structural models (D-E). Note the differences in HNH domain location, guide RNA direction, and gRNA/tDNA length. (F) Overlay of the fully-duplexed R-IscB-VR4 structure and the AlphaFold3-predicted R-IscB model. Residues beyond N434 could not be modeled in the density due to high mobility. (G) Representative BLI traces of R-IscB-VR4 and its M402A/D403A variant binding to an ssRNA substrate. (H) Representative association rate constants of R-IscB-VR4 RNP and its variants. A complete table of *k*_a_, *k*_dis_, and Kd is provided in Table S2.

### Structure-inspired residue substitutions further improve ssRNA binding in R-IscB

Next, we sought to further improve the ssNA-binding activity of R-IscB based on the new mechanistic insights. The structures reveal that a P1D loop extends outward like a lip, straining the base-pairing geometry at the beginning of the seed duplex. We therefore introduced three mutation clusters to either reduce the side-chain bulk of residues in this P1D loop or shorten its length by one amino acid. Among these, the R-IscB M402A/D403A double substitution was found to bind ssRNA targets 40-fold faster in *k*_on_ and 22-fold tighter in K_d_ than the parental version (Figure 4G), as measured by biolayer interferometry (BLI). We consider the association rate constant a more reliable indicator of R-IscB performance, as *k*_off_ is too slow to be quantified with confidence. Our structural analysis further revealed that even after 20 minutes of incubation at 37°C, approximately 50% of IscB remains trapped in the seed-duplexed state, suggesting that eviction of the HNH roadblock is a rate-limiting step in target recognition. Lowering the energetic barrier for HNH roadblock eviction would, in theory, accelerate target recognition and lead to more robust RNA-targeting performance in vivo. Our structures reveal that F196 stabilizes HNH in the roadblocked conformation by inserting into a hydrophobic pocket at the HNH-RuvC interface. We therefore designed two substitutions, F196H and F196W, with the goal of weakening this ball-and-socket interaction. F196H was found to bind ssRNA targets 2-fold faster in *k*_on_ and 3-fold tighter in K_d_ than the parental version, supporting our mechanistic hypothesis. As a consequence of TID removal, the linker from P1D to TID is absent from the EM density, and the linker connecting RuvC to P1D shows signs of elevated flexibility. To explore whether this elevated conformational flexibility might contribute to the weakened density observed at the first nucleotide of the gRNA-tRNA duplex, we attempted to remove one amino acid from the RuvC-P1D linker. The resulting variant (C378HAKN382 to AKIN) showed slightly improved binding behavior relative to the parental version (Figure 4H). Additional tested designs that were either neutral or detrimental to ssRNA binding are documented in Table S2 and Figure S7.

## Discussion

In this study, we attempted to provide the structural basis for single-stranded nucleic acid recognition in the context of full-length IscB and its truncated ssNA-targeting variant, R-IscB. It is reassuring that removal of the TAM-interaction domain does not alter the overall structure or the RNA-guided target-recognition mechanism. Based on these structures, we conclude that ssNA binding is a natural function of wild-type IscB, as elaborate conformational switching mechanisms are built in to ensure efficient target discovery through seed-duplex formation and stringent target validation through the seed-to-full duplex transition. We consider the TID a gain-of-function domain that redirects IscB’s activity toward dsDNA targets, as constant TAM-searching by the TID interferes with ssNA binding.

We reveal the presence of an elaborate conformational switching mechanism that serves two lines of important functions. It functions as an autoinhibition mechanism to downregulate the nuclease activity at the resting state, preventing IscB from non-specific sequence targeting of random nucleic acids. Such activities could be quite toxic to the host cell. The blockage of RuvC active site by the guide RNA is particularly noteworthy as it is simple yet efficient. The conformational switch also serves as a quality control mechanism to prevent off-targeting on a partial-matching target. Base-pairing beyond the seed region is required to pay the energetic cost of opening the HNH gate. We note that these two mechanisms are conserved in principle in larger Cas9 RNPs, even though the specific molecular contacts are not. Early R-loop states were captured by cryo-EM for SpCas9^29^, in which the authors show alternatively parked HNH and occluded HNH active site. The RuvC active site has not been observed to be blocked by the gRNA, however, multiple domain insertions trap the RuvC active site so deep inside that it may be sufficiently sequestered that way. Overall, our work reenforces the idea that Cas enzymes are conditional nucleases activated not only by RNA-guided substrate recruitment, but also by autoinhibition in the resting state and conditional activation in the target-bound state. We have observed at least two types of conditional activation mechanism, either an allosteric activation that controls the accessibility of the nuclease active site^30^ or a conditional recruitment mechanism of the nuclease *in trans*^*31*^.

Focusing on the effort to better use the utility of R-IscB, we have gained mechanistic insights from the comparison of full-length and R-IscB structures. Structural features straining the binding of ssNA have been identified. The realization of a steep kinetic and energetic barrier present in the target recognition process is instrumental. Initial efforts to incorporate these insights into further engineering has produced positive leads, which will be further tested in in vivo RNA editing experiments. Even though we focused this study on ssNA binding, we anticipate the majority of the mechanistic insights will be applicable to the dsDNA recognition process.

## Acknowledgements

This work was supported by the National Institutes of Health (NIH) under grant number R35GM118174 to A.K.

## METHOD DETAILS

### Plasmid construction

Engineered OgeuIscB and R-IscB-VR4 plasmids for RNP purification have been described previously^5,28^. ωRNA-V2 uses the optimized scaffold sequence and has a 16-nt guide at the 5’ side (5’-AAAAGAGTGAACGAGA). ωRNA was cloned into pUC57, in between HindIII and EcoRI, and under the control of an lpp promoter and a T7 terminator. Mutations in R-IscB were introduced into the R-IscB-VR4 expression vector by PCR-based site-directed mutagenesis. For in vivo editing experiments, codon-optimized R-IscB-VR4 plasmid has been described previously. Mutations and guide sequences were introduced via PCR mutagenesis. See Supplementary Tables for key sequence information.

### Protein expression and purification

Plasmids encoding IscB (enOgeuIscB, R-IscB-VR4 and its variants) and ωRNA were co-transformed into E. coli T7 Express plysY cells. The cell culture was grown in an LB medium supplemented with 0.75g L-cysteine/L at 37°C until the optical density at 600 nm reached 0.8. Expression was induced by adding isopropyl-β-D-thiogalactopyranoside (IPTG) to a final concentration of 0.5 mM at 16°C overnight. Cells were collected by centrifugation and lysed by French press in buffer A (50 mM HEPES pH 7.5, 300 mM NaCl, 0.5 mM TCEP, 5% glycerol, 2.5 mM MgCl_2_) with 1 mM phenylmethylsulfonyl fluoride (PMSF). The supernatant after centrifugation was applied onto strep-tactin XT resin (IBA lifesciences) preequilibrated in buffer A. Resin was then washed with 15mL of buffer A plus 0.1 mM CaCl_2_ and 10U DNase I (Thermo Scientific), 20mL of buffer B (50 mM HEPES pH 7.5, 1 M NaCl, 0.5 mM TCEP, 2.5 mM MgCl_2_), and 40mL of buffer A. Proteins were eluted from the resin using 10 mL of buffer A plus 50mM Biotin. The IscB-containing fractions were pooled, desalted, and further purified on MonoQ 5/50GL (Cytiva) with a NaCl gradient. The desired IscB fractions were pooled, concentrated, and further purified by size-exclusion chromatography (Superdex 200 Increase 10/300 GL; Cytiva) equilibrated with buffer C (50 mM HEPES pH 7.5, 150 mM NaCl, 0.5 mM TCEP, 2.5 mM MgCl_2_). The first peak was collected, concentrated, and snap-frozen in liquid nitrogen and stored at -80 °C.

### Biolayer interferometry (BLI)

BLI tests were conducted on Octet R8 (Satorius) in a BLI buffer of 50mM HEPES pH7.5, 40mM NaCl, 2.5 mM MgCl_2_, 0.05% Tween-20, and 0.1% BSA at 37 °C under a shaking speed of 1,000 rpm. Briefly, streptavidin biosensors (Satorius) were first loaded with ssRNA (500 nM biotinylated ssRNA in BLI buffer), baselined, associated in defined concentrations of R-IscB variants (10, 30, 100, 300 nM in BLI buffer), and dissociated in the buffer with no proteins. Baseline correction was conducted in Octet Analysis Studio v13.0.3.52 software, and the corrected traces were plotted and analyzed using GraphPad Prism.

### Cryo-EM sample preparation, data acquisition, and processing

DNA oligonucleotides for cryo-EM were synthesized (Integrated DNA Technologies). IscB was incubated for 20 minutes at 37°C with the target ssDNA/RNA substrates in binding buffer (50 mM HEPES pH 7.5, 50 mM NaCl, 2 mM DTT, 2.5 mM MgCl_2_). Substrates were supplied at a 2-fold molar excess to IscB (0.5mg/mL final concentration). 3.5μL of were applied to a Quantifoil holey carbon grid (1.2/1.3, Cu 200 mesh) which had been glow-discharged with 25mA at 0.39 mBar for 30 seconds (PELCO easiGlow). Grids were blotted with Vitrobot blotting paper (Electron Microscopy Sciences) for 5.5 s at 4 °C, 100% humidity, and plunge-frozen in liquid ethane using a Mark IV Vitrobot (Thermo Fisher). Data were collected on a Krios G3i Cryo Transmission Electron Microscope (Thermo Scientific) with a Ceta 16M CMOS camera 300kV, Gatan K3 direct electron detector. The total exposure time of each movie stack led to a total accumulated dose of 50 electrons per Å^2^ which fractionated into 40 frames. Dose-fractionated super-resolution movie stacks collected from the Gatan K3 direct electron detector were collected at a pixel size of 0.4125 Å. The defocus value was set between −1.0 μm to −2.5 μm.

Motion correction, CTF-estimation, blob particle picking, 2D classification, 3D classification and non-uniform 3D refinement were performed in cryoSPARC v.4.7.1^32^. Refinements followed the standard procedure, a series of 2D and 3D classifications with C1 symmetry were performed as shown in Fig. S2-3 to generate the final maps. A solvent mask was generated and was used for all subsequent local refinement steps. CTF post refinement was conducted to refine the beam-induced motion of the particle set, resulting in the final maps. The detailed data processing and refinement statistics for cryo-EM structures are summarized in Fig. S2-3 and Table S1.

---

**Supplementary Figure 1.**
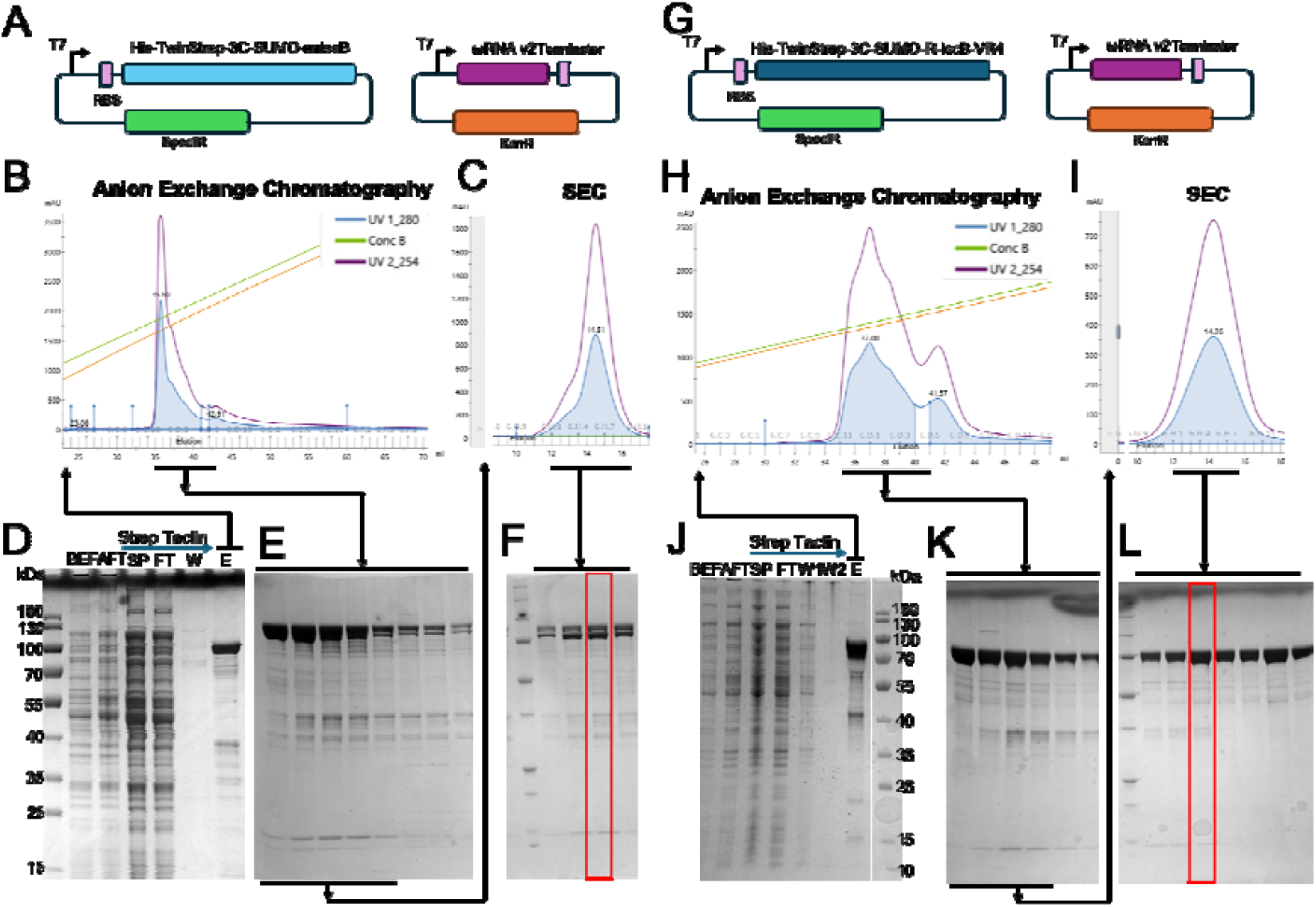
Reconstitution of OgeuIscB-ωRNA RNPs. (A-F) Workflow of engineered IscB RNP purification. (A) Co-expression scheme of enIscB and ωRNA plasmids. (B) Elution profile of the enIscB-ωRNA RNP on anion exchange chromatography. (C) Elution profile of enIscB-ωRNA RNP on size-exclusion chromatography (SEC). (D) SDS-PAGE analysis of the Strep-tactin purified enIscB-ωRNA RNP. Before induction (BEF), after induction (AFT), lysate supernatant (SP), strep resin flow thru (FT), wash(W), and elution (E). (E) SDS-PAGE of anion exchange peak fractions. (F) SDS-PAGE of SEC peak fractions. (G-M) Workflow of R-IscB-VR4 RNP purification. (G) Co-expression scheme of R-IscB-VR4 and ωRNA plasmids. (H) Elution profile of the R-IscB-VR4-ωRNA RNP on anion exchange chromatography. (I) Elution profile of R-IscB-VR4-ωRNA RNP on size-exclusion chromatography (SEC). (J) SDS-PAGE analysis of the Strep-tactin purified R-IscB-VR4-ωRNA RNP. Before induction (BEF), after induction (AFT), lysate supernatant (SP), strep resin flow thru (FT), Dnase I wash (W1), wash2 (W2), and elution (E). (K) SDS-PAGE of anion exchange peak fractions. (L) SDS-PAGE of SEC peak fractions.

**Supplementary Figure 2.**
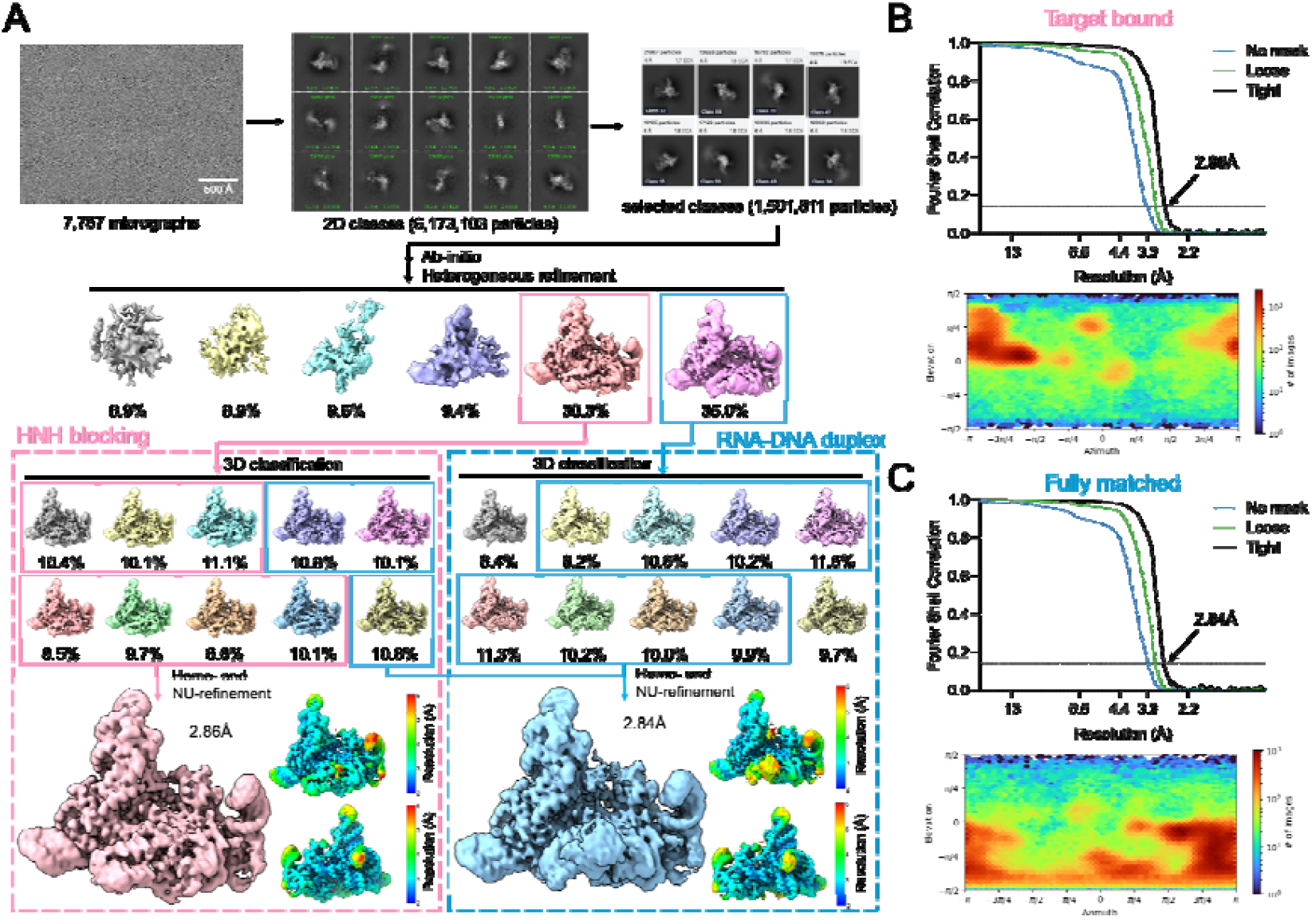
Cryo-EM single particle reconstruction of enIscB-ωRNA-ssDNA complex. (A) Workflow of the cryo-EM image processing and 3D reconstruction for the enIscB-ωRNA-ssDNA complex. 3D classification was used to separate and reassign the particles to two states. (B-C) Top: Fourier Shell Correlations (FSC) of enIscB-ωRNA-ssDNA complex reconstruction with the gold-standard cutoff (FSC = 0.143) marked with a dotted line for (B) target bound state and (C) fully matched state. Bottom: Direction distribution plot for each state, respectively.

**Supplementary Figure 3.**
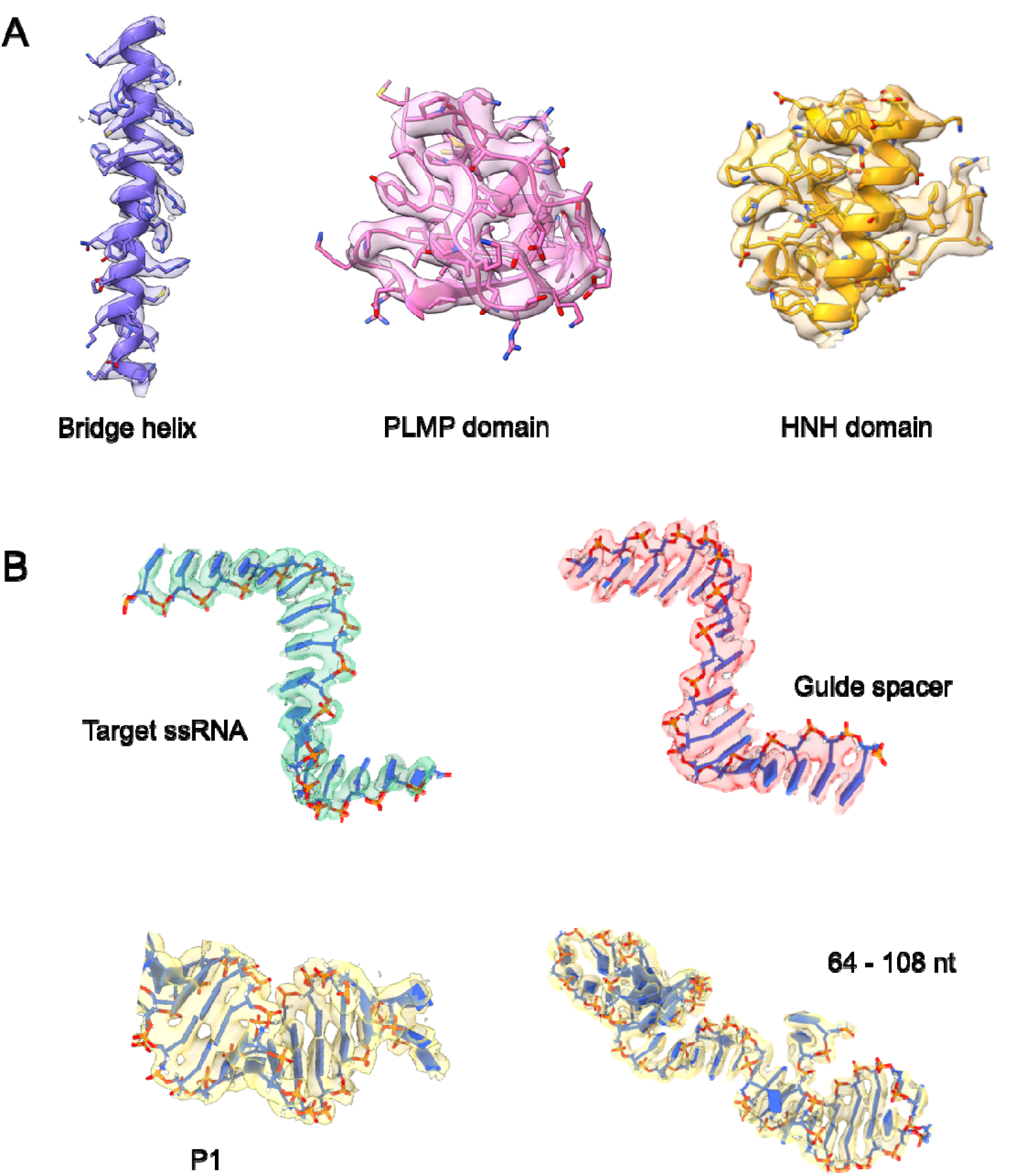
Representative local map density for the different functional states. (A) EM densities for representative protein regions inside R-IscB-VR4-ωRNA-ssRNA complex. (B) EM densities for the target ssRNA strand and RNA regions inside the R-IscB-VR4-ωRNA-ssRNA complex.

**Supplementary Figure 4.**
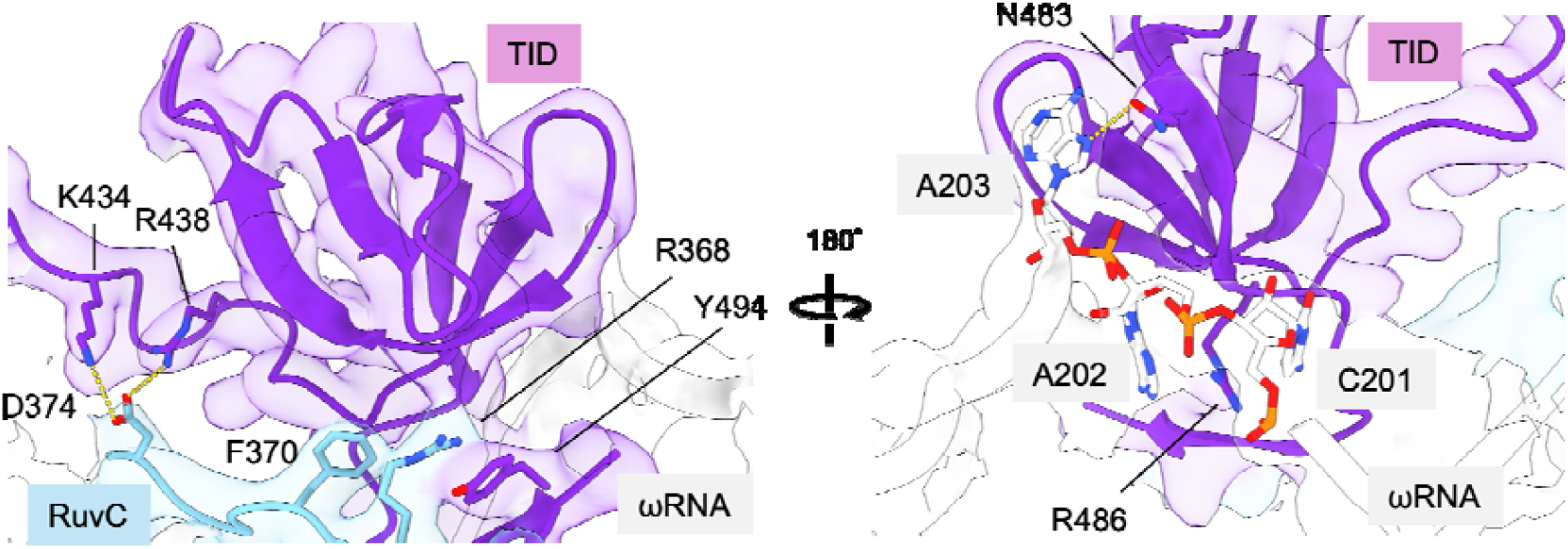
Zoom-in views show the interactions between TID and adjacent RuvC domain and ωRNA, including salt bridge, cation-π stacking and hydrogen bonds.

**Supplementary Figure 5.**
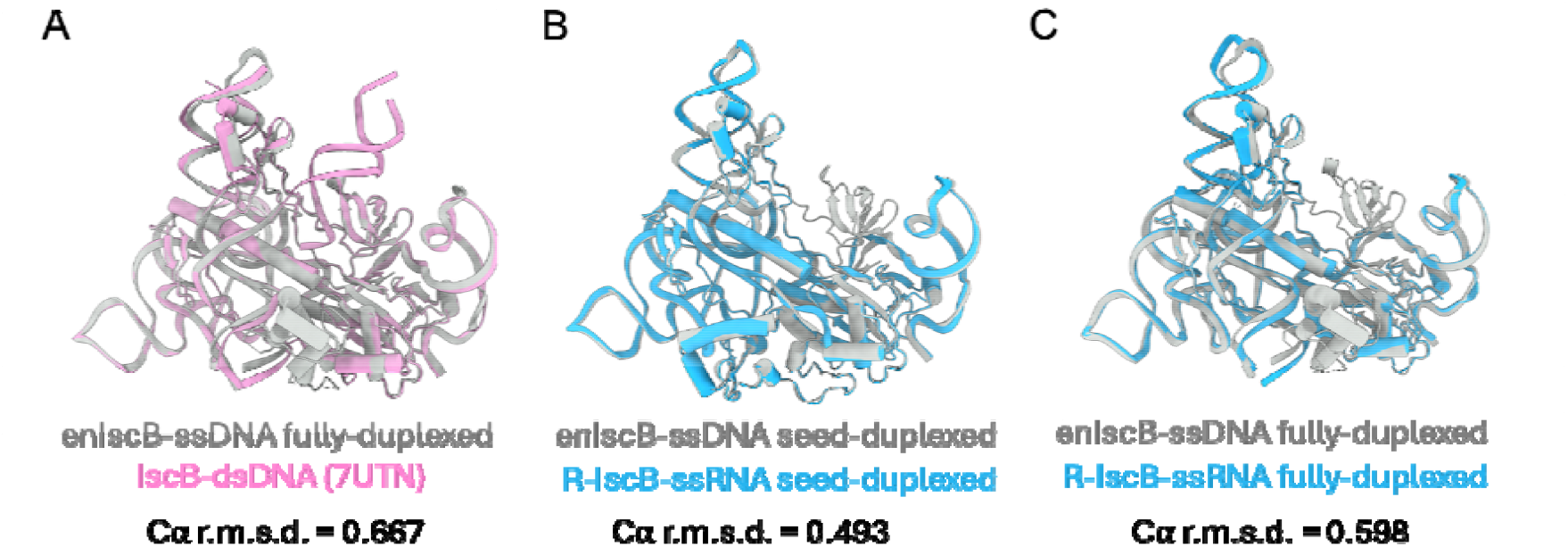
Structure comparison of IscB and its variants. (A) Overlay of enIscB-ssDNA fully-duplexed state and previously reported IscB-dsDNA (PDB: 7UTN). (B) Overlay of enIscB-ssDNA and R-IscB-VR4-ssRNA in the seed-duplexed state. (C) Overlay of enIscB-ssDNA and R-IscB-VR4-ssRNA in the fully-duplexed state.

**Supplementary Figure 6.**
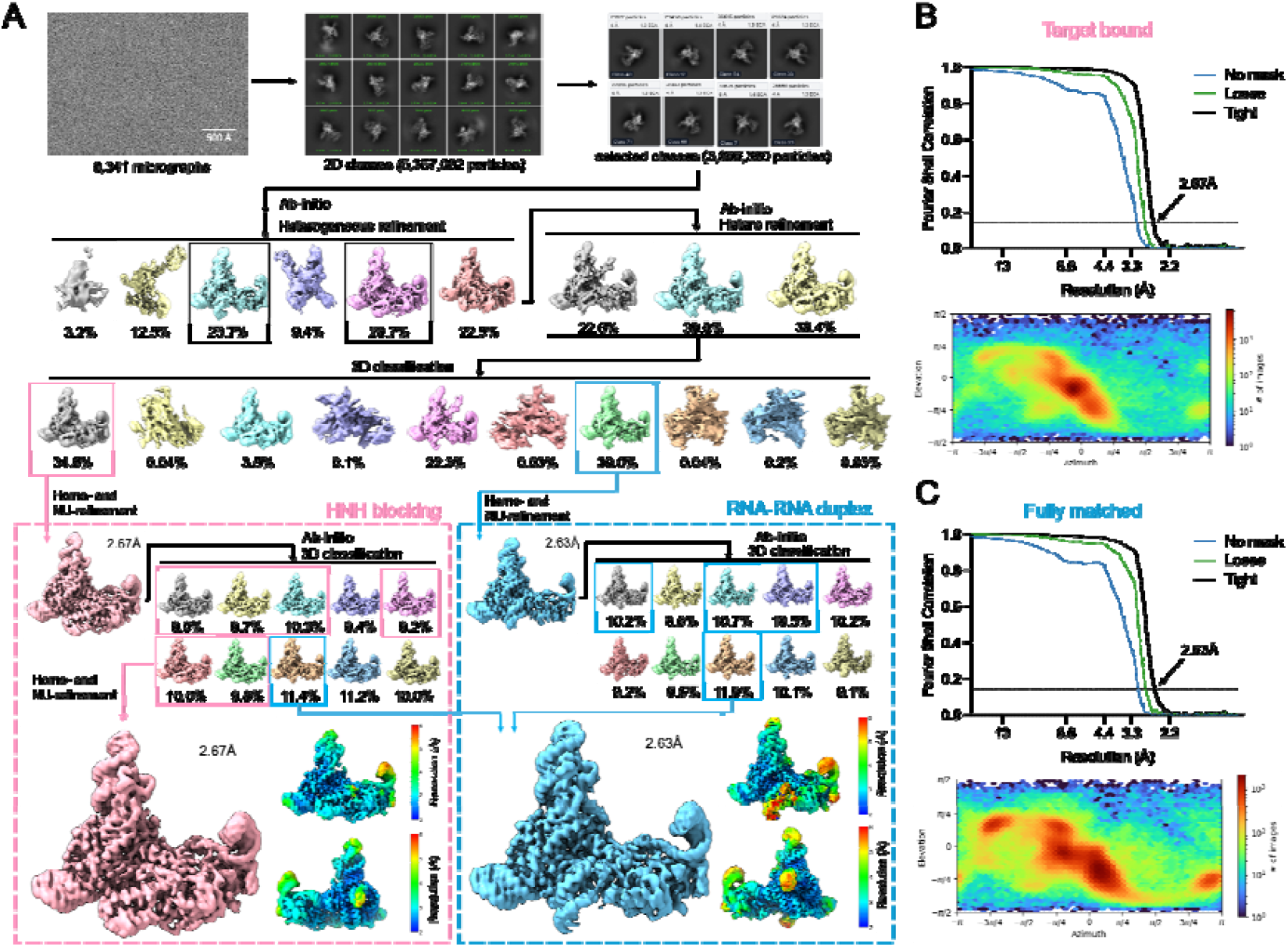
Cryo-EM single particle reconstruction of R-IscB-VR4-ωRNA-ssRNA complex. (A) Workflow of the cryo-EM image processing and 3D reconstruction for the R-IscB-VR4-ωRNA-ssRNA complex. 3D classification was used to separate and reassign the particles to two states. (B-C) Top: Fourier Shell Correlations (FSC) of R-IscB-VR4-ωRNA-ssRNA complex reconstruction with the gold-standard cutoff (FSC = 0.143) marked with a dotted line for (B) target bound state and (C) fully matched state. Bottom: Direction distribution plot for each state, respectively.

**Supplementary Figure 7.**
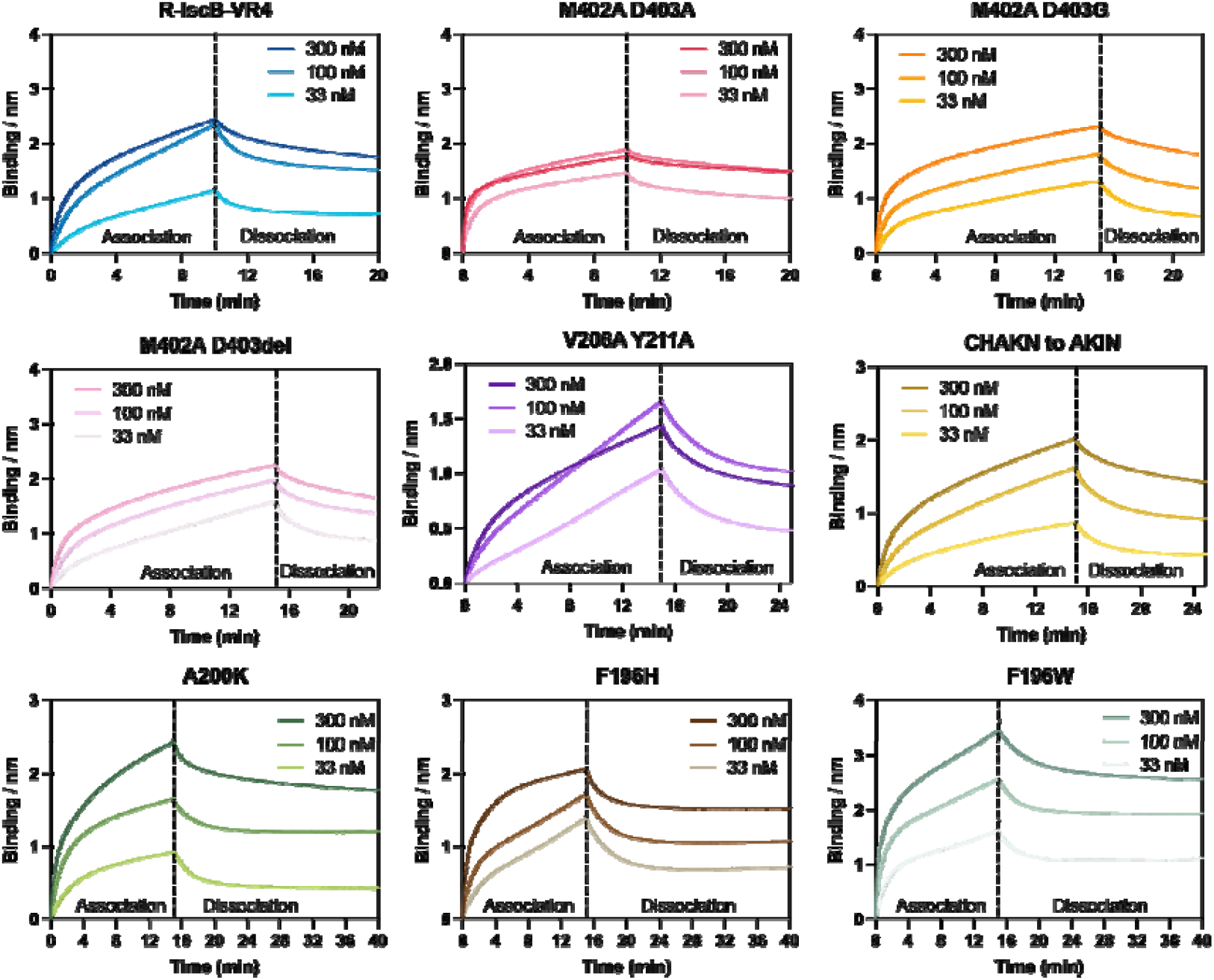
Biolayer interferometry results of R-IscB-VR4 and its variants. Biotinylated ssRNA substrate was loaded onto streptavidin biosensor tips. Binding is represented by wavelength shift (Δλ, measured in nanometers) measured by Octet instrument.

**Table S1.**
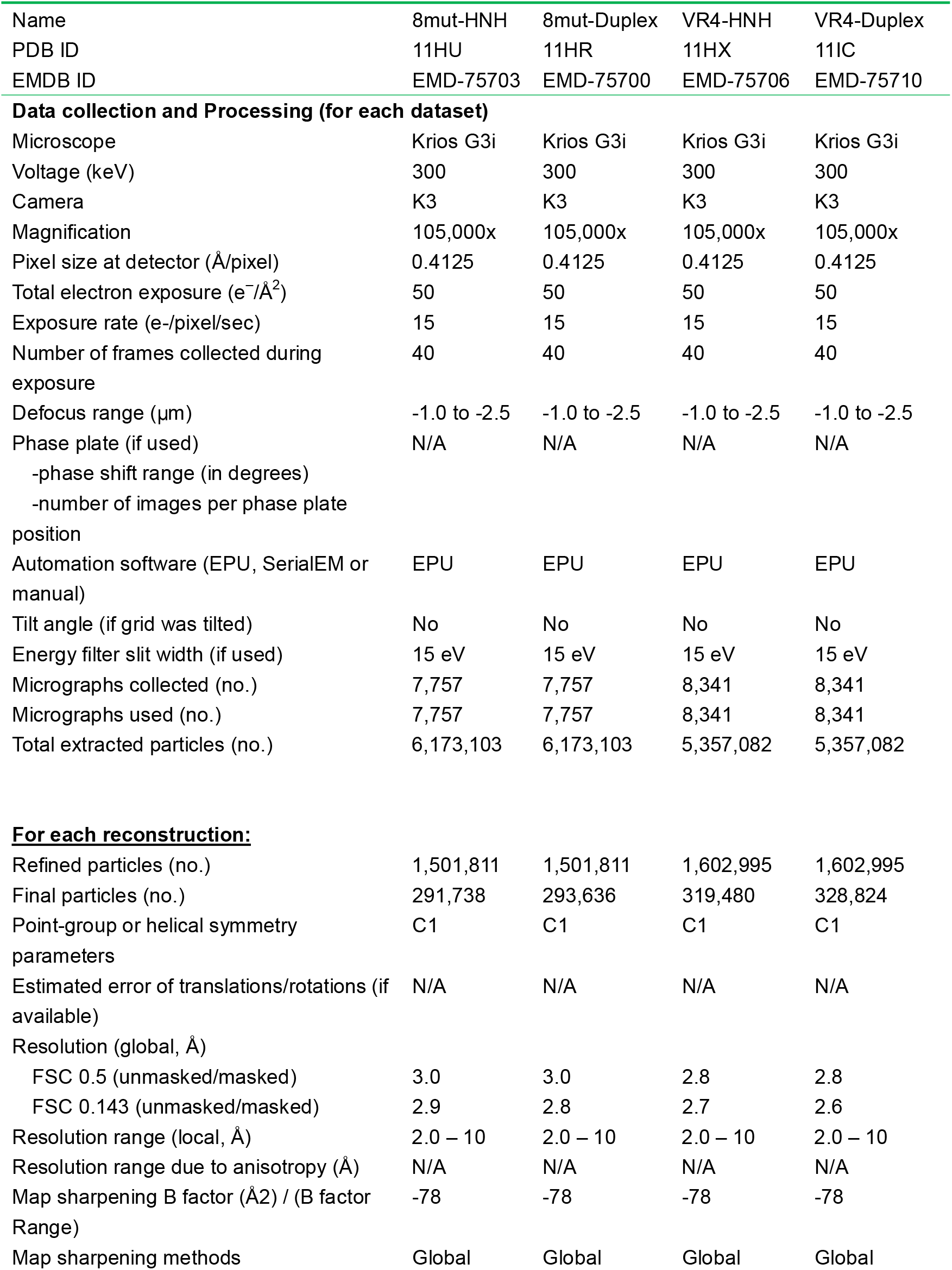
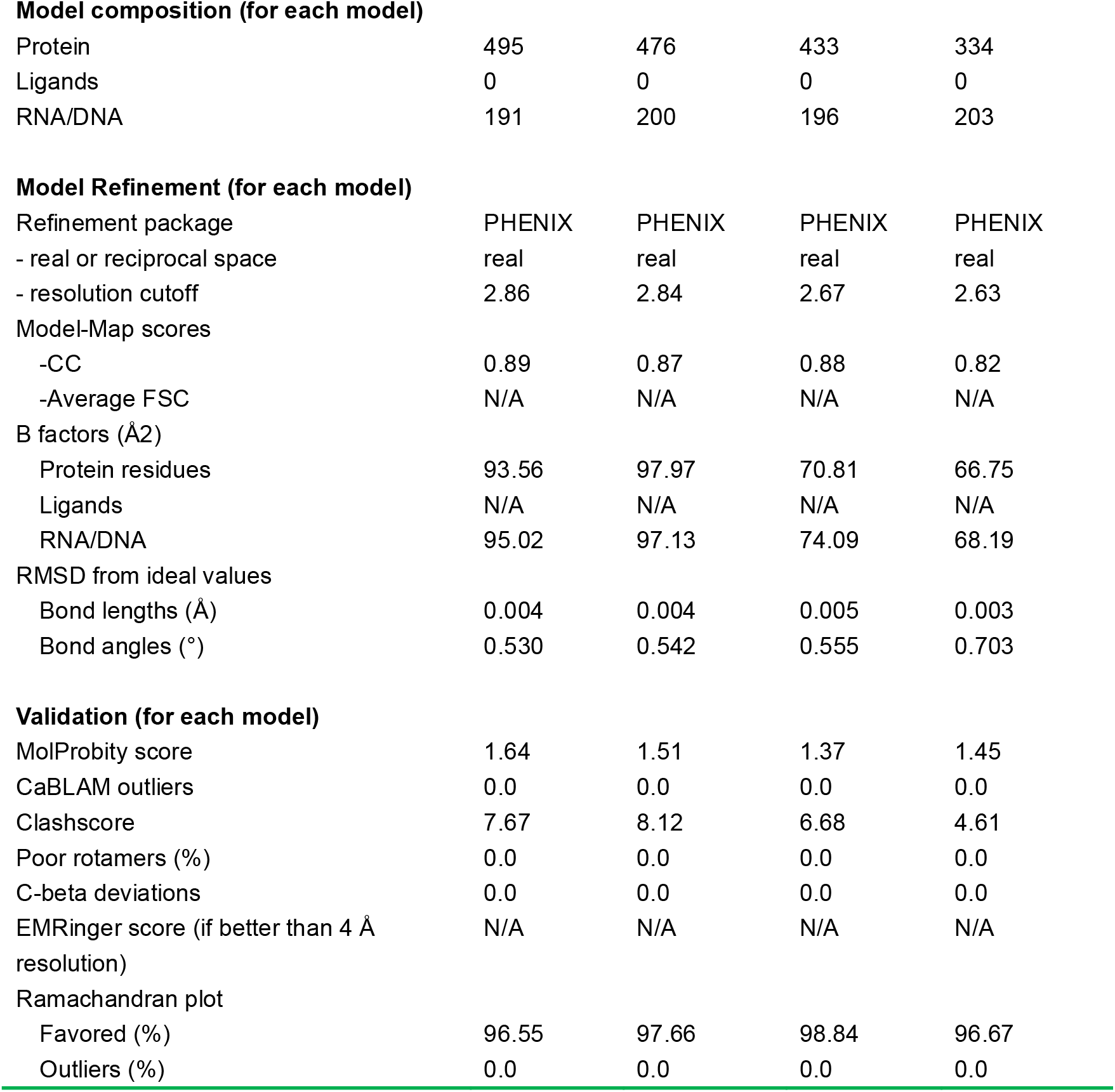
Cryo-EM data collection, refinement, and validation statistics.

**Table S2.**
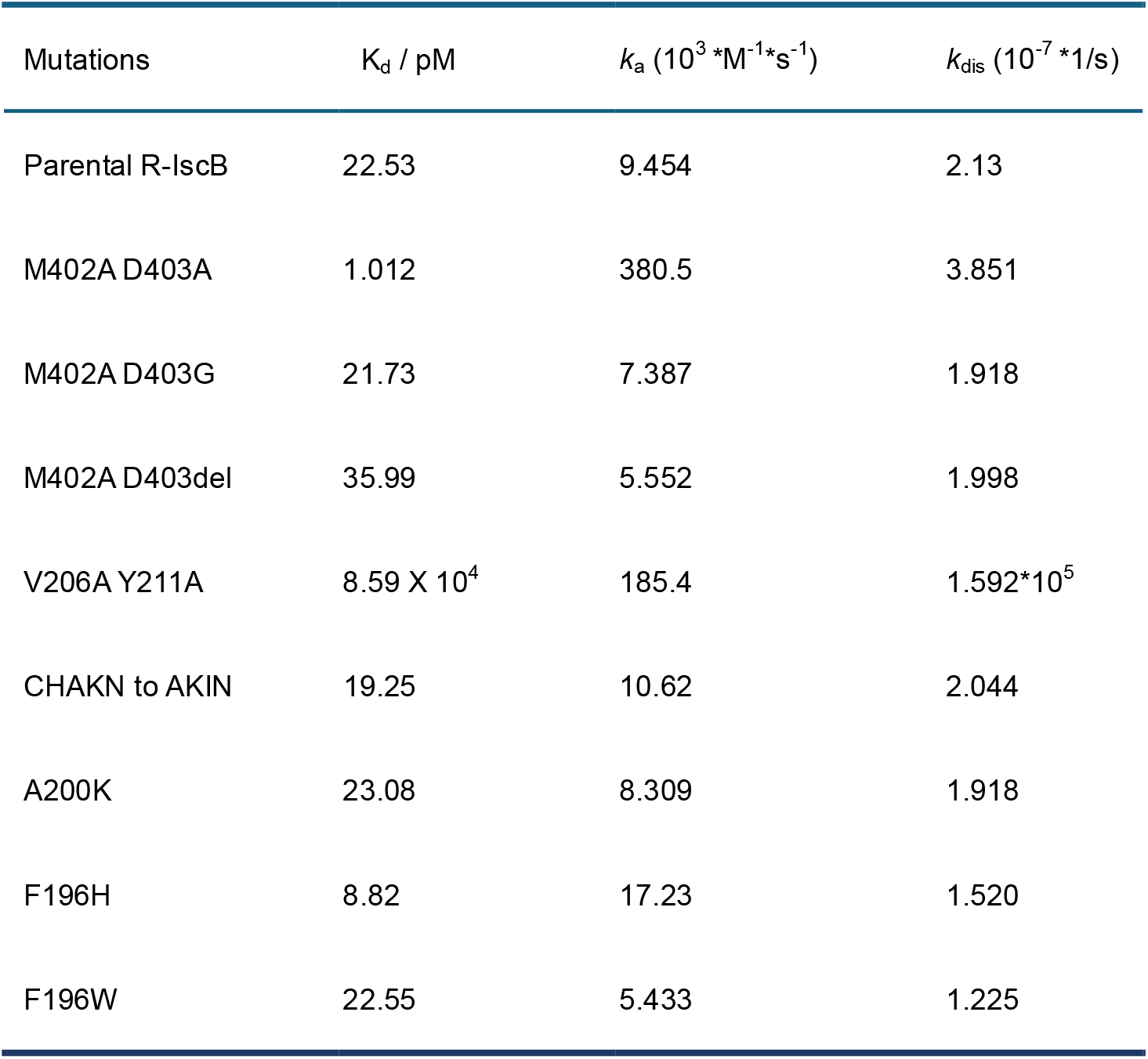
Kinetic measurements of R-IscB-VR4 RNP and its variants.

| Mutations | $K_d$ / pM | $k_a$ ($10^3 \cdot \text{M}^{-1} \cdot \text{s}^{-1}$) | $k_{\text{dis}}$ ($10^{-7} \cdot \text{s}^{-1}$) |
| --- | --- | --- | --- |
| Parental R-IscB | 22.53 | 9.454 | 2.13 |
| M402A D403A | 1.012 | 380.5 | 3.851 |
| M402A D403G | 21.73 | 7.387 | 1.918 |
| M402A D403del | 35.99 | 5.552 | 1.998 |
| V206A Y211A | $8.59 \times 10^4$ | 185.4 | $1.592 \cdot 10^5$ |
| CHAKN to AKIN | 19.25 | 10.62 | 2.044 |
| A200K | 23.08 | 8.309 | 1.918 |
| F196H | 8.82 | 17.23 | 1.520 |
| F196W | 22.55 | 5.433 | 1.225 |

